# Genetic bioaugmentation-mediated bioremediation of terephthalate in soil microcosms using an engineered environmental plasmid

**DOI:** 10.1101/2024.08.19.608593

**Authors:** Alejandro Marquiegui Alvaro, Anastasia Kottara, Micaela Chacón, Lisa Cliffe, Michael Brockhurst, Neil Dixon

## Abstract

Harnessing in situ microbial communities to clean-up polluted natural environments is a potentially efficient means of bioremediation, but often the necessary genes to breakdown pollutants are missing. Genetic bioaugmentation, whereby the required genes are delivered to resident bacteria via horizonal gene transfer, offers a promising solution to this problem. Here we engineered a conjugative plasmid previously isolated from soil, pQBR57, to carry a synthetic set of genes allowing bacteria to consume terephthalate, a chemical component of plastics commonly released during their manufacture and breakdown. Our engineered plasmid caused a low fitness cost and was stably maintained in terephthalate contaminated soil by the bacterium *P. putida.* Plasmid carriers efficiently bioremediated contaminated soil, achieving complete breakdown of 3.2 mg/g of terephthalate within 8 days. The engineered plasmid horizontally transferred the synthetic operon to *P. fluorescens in situ*, and the resulting transconjugants degraded 10 mM terephthalate during a 180-hour incubation. Our findings show that environmental plasmids carrying synthetic catabolic operons can be useful tools for *in situ* engineering of microbial communities to perform clean-up even of complex environments like soil.

## Introduction

As global plastic production continues to rise, so does the demand of raw materials for plastic manufacture. Polyethylene terephthalate (PET) plastic has an annual global production that surpasses 85 million metric tons, and is ubiquitous in packaging and textile industries ^1^. Alongside other polyester-based plastics such as polybutylene terephthalate (PBT) used as a thermal insulator ^2^, PET is comprised of terephthalate (TPA) monomers, in addition to ethylene glycol (EG). The uses of TPA are not limited to polymer synthesis, as it is also used as an elastomer and plasticizer to enhance material formulation and properties ^3,4^. The chemical production of TPA requires a purification step that yields purified TPA and wastewater, and this wastewater is contaminated with a range of aromatic hydrocarbons, primarily TPA with concentrations as high as 500 mg/L^5^. The release of untreated TPA-contaminated wastewater into soil ecosystems poses an environmental threat due to its toxicity ^6,7^. Additionally, TPA release can arise from *in situ* biodegradation of TPA-containing polyester plastics by microorganisms ^8–11^.

Microbial biodegradation and consumption of TPA offers a promising potential solution to TPA contamination. Bacteria such as *Comamonas* sp. E6, *Rhodococcus jostii* RHA1or *Pseudomonas umsongensis* sp. G016 have all been reported to degrade and consume TPA ^12–14^ and TPA-degrading operons have been found bioinformatically across diverse bacterial taxa ^15,16^. Use of an anaerobic microbial community to treat TPA contaminated wastewater have shown effective decontamination ^17^. However, whether such microbial bioremediation of TPA would work in more complex natural environments, such as soils, remains unclear.

The addition of exogenous microorganism(s) to a natural environment, in order to introduce or modify existing community traits, referred to as bioaugmentation, can be effective for a range of applications in soil environments including nitrogen fixation ^18,19^, phosphorous bioavailability ^20, 21^ and bioremediation of organic pollutants ^22–25^ or heavy metals ^26,27^. Notably, bioaugmentation for bioremediation can enable clean-up of already contaminated environments (*in situ* bioremediation). Compared to physicochemical methods for pollutant removal (e.g. soil washing, incineration, chemical oxidation), bioaugmentation offers a cheaper and less disruptive alternative ^28^. Two main types of bioaugmentation for bioremediation exist: cellular and genetic bioaugmentation. The former introduces the bioremediation trait within a focal microorganism that itself then performs the breakdown of the pollutant, while the latter introduces the genes for the bioremediation trait encoded upon a mobile genetic element within a donor microorganism which can both perform breakdown and transfer the bioremediation trait to neighbouring cells (recipients) (**Figure 1A**). The resident native microbiota in any given natural environment is likely to be well-adapted and occupy multiple niches providing high colonization resistance ^29^. As such, introduced exogenous microorganisms frequently experience low survival ^23,30–32^. It has been hypothesized, therefore, that genetic bioaugmentation may be more effective than cellular bioaugmentation in natural communities because it does not rely upon long-term persistence of the exogenous microbe provided it is resident long enough to transfer the mobile genetic element to resident microorganism(s)^28^. Indeed, experimental evidence of stable maintenance of the pollutant-degrading genes upon a mobile element despite rapid extinction of the donor microorganism has been reported ^33–36^.

**Figure 1.**
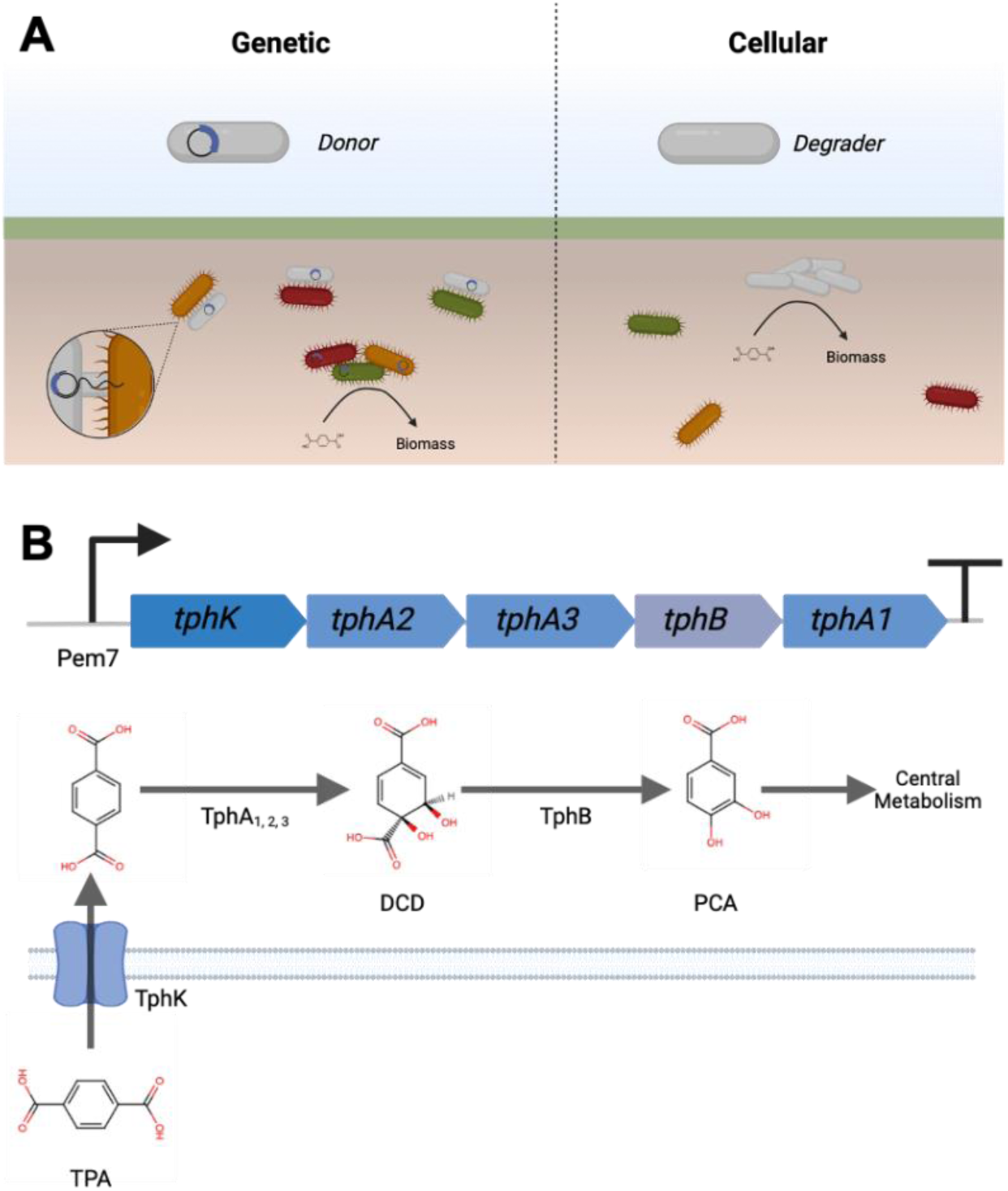
**(A)** Left panel represents genetic bioaugmentation where upon donor inoculation the plasmid carrying the catabolic genes is horizontally transferred (via conjugation) to the resident members of the community. Upon acquiring the catabolic genes, the resident microbiome transforms the pollutant to biomass thus achieving bioremediation. Right panel represents cellular bioaugmentation where the inoculated bacterium is responsible for the conversion of pollutant into biomass. (**B**) Top, TPA-degrading operon architecture; bottom, TPA biodegradation pathway: TPA is imported into the cell via *tphK* where it is converted to 1,2-dihydroxy-3,5-cyclohexadiene-1,4-dicarboxylate (DCD) by *tphA_1_A_2_A_3_*and then into protocatechuic acid (PCA) by *tphB*. PCA enters central metabolism via the Krebs cycle.

Genetic bioaugmentation mimics the natural evolutionary process of horizontal gene transfer (HGT), which enables rapid adaptation by microorganisms to environmental fluctuations through acquisition of novel traits from neighbouring cells^37^, including those for dealing with pollutant accumulation in soil^38^ and catabolic operons^39–41^. Like natural HGT, genetic bioaugmentation has taken advantage of diverse mobile genetic elements for transferring traits, including phage transduction and plasmid conjugation. Conjugative plasmids, equipped with the machinery required for conjugation (*tra*, *mob* and *oriT*), are particularly attractive tools for genetic bioaugmentation because they can carry large genetic cargos, including multiple entire multi-gene operons. For instance, the TOL plasmid pWW0 contains the catabolic operon for toluene and xylene utilization^42^, and the NAH plasmid is equipped with naphthalene-degrading enzymes^43^. Successful implementation of genetic bioaugmentation using conjugative plasmids has been reported across a range of plasmids, donors, microbial communities, and terrestrial environments^22,23,33,44,45^.

Engineering natural plasmid backbones isolated from the target environment is likely to have several advantages over using model plasmids typically used in molecular biology. Such advantages include having been previously selected in nature to conjugate efficiently in the environmental substrate which is likely to be more complex than lab media (e.g., soils) and having evolved host ranges that are suitable for spreading into common taxa within the resident microbiota. To promote effective genetic bioaugmentation for *in situ* pollutant degradation an ideal plasmid vector would have a high conjugation rate, to enable plasmid spread before the donor becomes extinct, and a minimal fitness cost, to prevent lowering the recipients’ fitness upon plasmid acquisition in the chosen environmental context, as well as a broad host range, ensuring wide dissemination of the new trait in the native resident community. pQBR57 is a natural mercury-resistance encoding megaplasmid (307 kbp) isolated from the sugar beet rhizosphere in Oxfordshire^46^, which can conjugate into diverse species of Pseudomonas^47^, a highly diverse genus of Gram negative bacteria common in soils and freshwater. pQBR57 has a high conjugation rate and minimal fitness cost in *Pseudomonas fluorescens*, enabling the plasmid to spread within soil communities through interspecies conjugation^48,49^. Crucially, pQBR57 is also amenable to genetic manipulation^50^, enabling the insertion of new genetic cargos.

Here we explore the potential of the environmental conjugative plasmid pQBR57 to act as a genetic bioaugmentation vector. Firstly, we engineered pQBR57 to encode a synthetic TPA-degrading operon, *tph*, and tested its functionality in soil microcosms using *Pseudomonas putida* as the donor species. The operon, herein referred to as **KAB** operon, is composed of a TPA transporter (*tpa**K***) from *Pseudomonas mandelli*, and terephthalate dioxygenases (*tph**A**2*, *tphA3*), reductase (*tphA1*) and dehydrogenase (*tph**B***) from *Comamonas* sp. E6^51,52^ (**Figure 1B**). We show that *P. putida* with pQBR57-KAB can grow on TPA in both liquid media and soil microcosms, and also confirm that this results in depletion of TPA *in situ*. Furthermore, we demonstrate conjugative transfer of pQBR57-KAB in soil microcosms from *P. putida* to *P. fluorescens* and confirm that a substantial fraction of the plasmid recipients gained the ability to grow on and degrade TPA. In summary, we demonstrate the utility of using an environmental conjugative plasmid encoding synthetic catabolic operon for genetic bioaugmentation in soil.

## Results

### Assembly and functional testing of pQBR57-KAB in liquid

To develop a TPA-degrading operon, we first assembled the genes corresponding to the synthetic KAB operon (*tpa**K*** from *P. mandelli*, and *tph**A**2*, *tphA3*, *tphA1* and *tph**B*** from *Comamonas* sp. E6) and placed it under the control of an average-strength constitutive promoter, Pem7^53^. The assembled operon was then inserted between genes pQBR57_0027 and pQBR57_0028 via *in vivo* homologous recombination in *P. fluorescens* harbouring pQBR57 (**Figure 2A**). The resulting pQBR57-KAB was transferred via conjugation to *P. putida*. *P. putida* pQBR57-KAB, along with *P. putida* pQBR57, was grown in M9 media with 10 mM TPA. The *P. putida* pQBR57-KAB strain was able to utilize TPA as its sole carbon source resulting in a growth curve marked by a prolonged lag-phase (approx. 60 hours); in contrast *P. putida* pQBR57 did not grow (**Figure 2B**). The concentration of TPA at the end of the growth assay is significantly lower in *P. putida* pQBR57-KAB than *P. putida* pQBR57 (**Figure 2C**, Wilcoxon test, W = 64, p < 0.001) where the mean TPA concentration of *P. putida* pQBR57 is 9.47 mM TPA (N = 4) and those with pQBR57-KAB is 0.0 mM TPA (N = 4).

**Figure 2.**
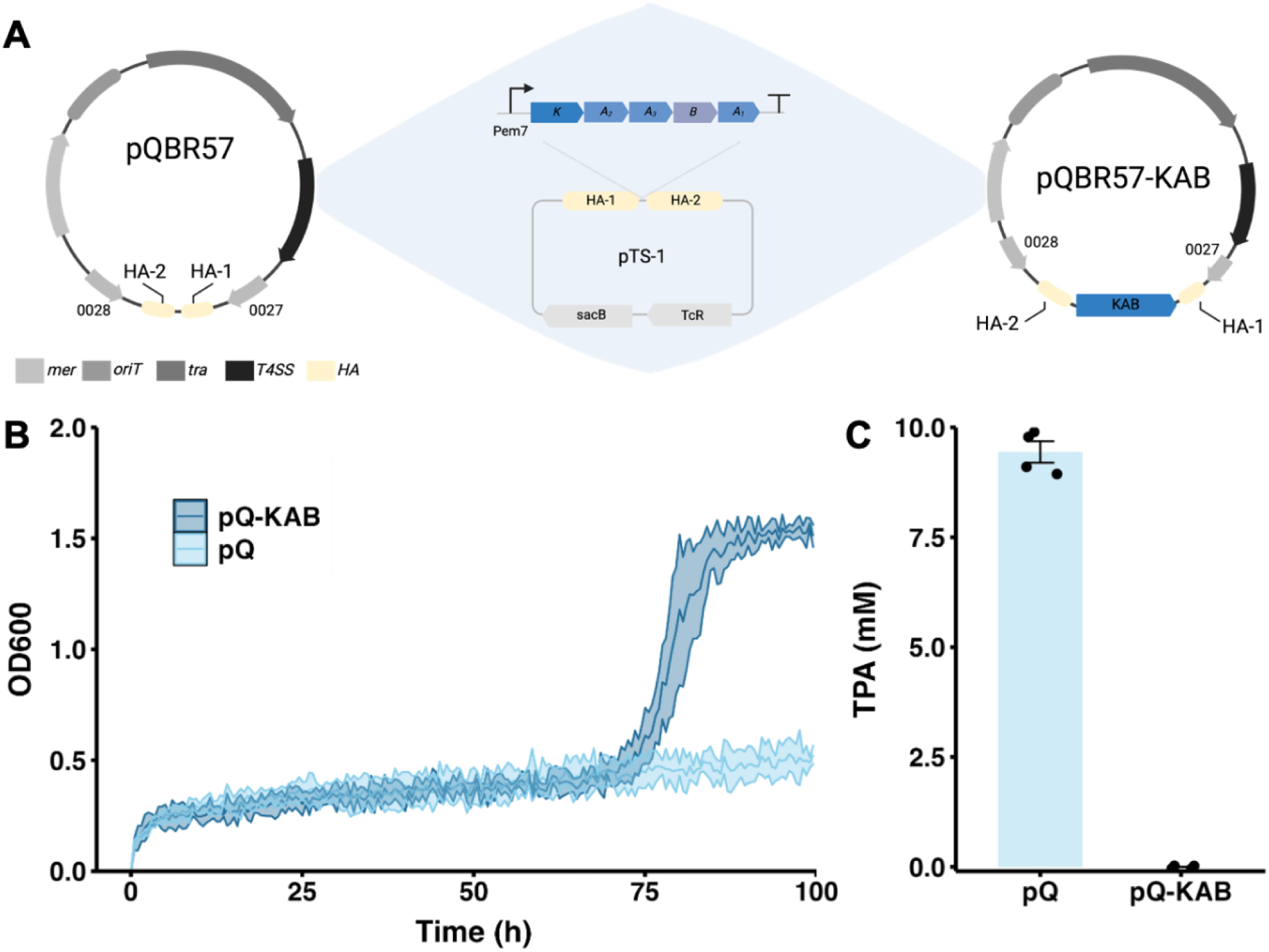
(**A**) Assembly of pQBR57-KAB. The original pQBR57 plasmid has mercury resistance (*mer*), as well as the determinant for conjugation including the origin of transfer (*oriT*), mating-pair formation (*tra*) and Type IV secretion system (*T4SS*). The homology arms (HA) are also found in the suicide vector pTS-1 to guide the homologous recombination of the KAB operon in the intergenic region between pQBR57_0027 and pQBR57_0028. The backbone of pTS-1 has tetracycline resistance (*TcR*) for selection and levensucrase (*sacB*) for counter-selection. On the right is the resulting pQBR57-KAB. (**B**) Growth dynamics of *P. putida* pQBR57 and pQBR57-KAB in M9 media with 10 mM TPA. (**C**) Final concentration of TPA after the growth dynamics experiment.

### Conjugation rate and fitness cost of pQBR57-KAB

To estimate the impact of the catabolic payload on plasmid transfer we measured the conjugation rate of pQBR57 and pQBR57-KAB by the Simonsen method^54^ using *P. putida* as the plasmid donor, and *P. putida* (intraspecies) or *P. fluorescens* (interspecies) as plasmid recipients. The conjugation rate depended on the combination of plasmid and recipient (**Figure 3A**, effect of plasmid by recipient interaction, factorial ANOVA, F_1,44_ = 5.19, p = 0.03). Whereas the intraspecific conjugation rates were equivalent for both plasmids, pQBR57-KAB had a lower interspecific conjugation rate than pQBR57 (**Figure 3A**). In addition, conjugation rates were higher when *P. fluorescens* is the recipient, regardless of plasmid genotype.

**Figure 3.**
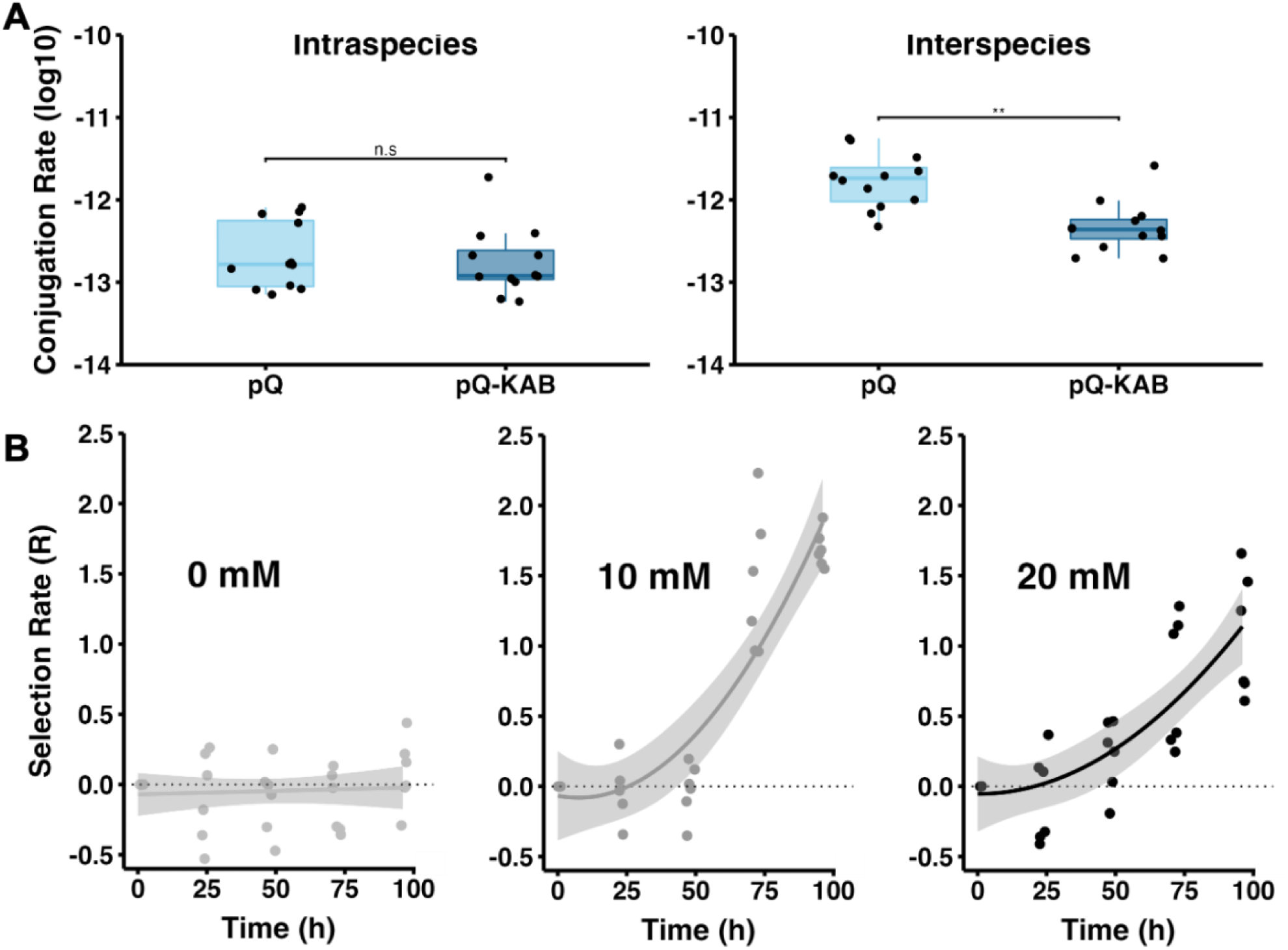
(**A**) The conjugation rate of pQBR57 and pQBR57-KAB, on the left is intraspecies (*P. putida* is donor and recipient) while on the right side is interspecies (*P. putida* is the donor and *P. fluorescens* is the recipient). The boxplots show the median conjugation rate (**B**) The selection rate of pQBR57-KAB compared to pQBR57 in *P. putida* in minimal media supplemented with 5 mM of glucose and a range of TPA concentrations (0, 10 and 20 mM).

The fitness effect of the KAB operon was analysed with competition assays between *P. putida* pQBR57-KAB and *P. putida* pQBR57. The strain carrying the KAB operon was chromosomally tagged with a fluorescent protein enabling direct quantification of each sub-population over time by flow cytometry. Each competition was carried out in M9 media supplemented with 5 mM glucose at a range of TPA concentrations (0, 10 and 20 mM). The fitness of *P. putida* pQBR57-KAB relative to *P. putida* pQBR57 increased with addition of TPA to the growth medium and was highest at intermediate concentrations (**Figure 3B**, effect of TPA by time interaction, repeated measures ANOVA, F_8,40_ = 50.09, p < 0.001). Selection for *P. putida* pQBR57-KAB was significantly stronger at 10 mM TPA than 20 mM TPA at both 72 and 96 hours (F_1,5_ = 34.81, p = 0.002, and F_1,5_ = 15.31, p = 0.01, respectively). Therefore, the KAB catabolic operon confers a significant fitness benefit in the presence of TPA, which is strongest at intermediate TPA concentrations. Moreover, the fitness cost of the catabolic payload is negligible in the absence of TPA, as the selection rate remains close to zero throughout the length of the experiment.

### TPA degradation and maintenance of pQBR57-KAB in soil microcosm

To assess the bioremediation function of pQBR57-KAB in soil, soil microcosms supplemented with 3.2 mg/g TPA were inoculated with *P. putida* pQBR57-KAB, *P. putida* pQBR57 or no bacteria. The concentration of TPA and population size were assessed after 3, 6, 8 and 32 days. Cumulative bacterial growth (measured by the area under the curve) increased only with TPA supplementation when bacteria carried pQBR57-KAB (**Figure 4A and 4B**, two-way ANOVA interaction effect, F_1,20_ = 97.36, p < 0.001). Whereas no appreciable TPA breakdown was observed within soil microcosms containing *P. putida* pQBR57 relative to soil without bacteria (**Supplementary Figure 3** Welch *t*-test, t_5.03_ = 0.0017, p = 0.99), by day 8 all the TPA had been consumed by *P. putida* pQBR57-KAB (**Figure 4C**, Wilcoxon test, W = 0, p = 0.003). Consistent with a strong fitness benefit of pQBR57-KAB in the presence of TPA, pQBR57-KAB was maintained at higher frequency with versus without TPA, whereas the frequency of pQBR57 was unaffected by TPA (**Figure 4D**, two-way ANOVA effect of plasmid by TPA, F_1,20_ = 13.50, p = 0.002).

**Figure 4.**
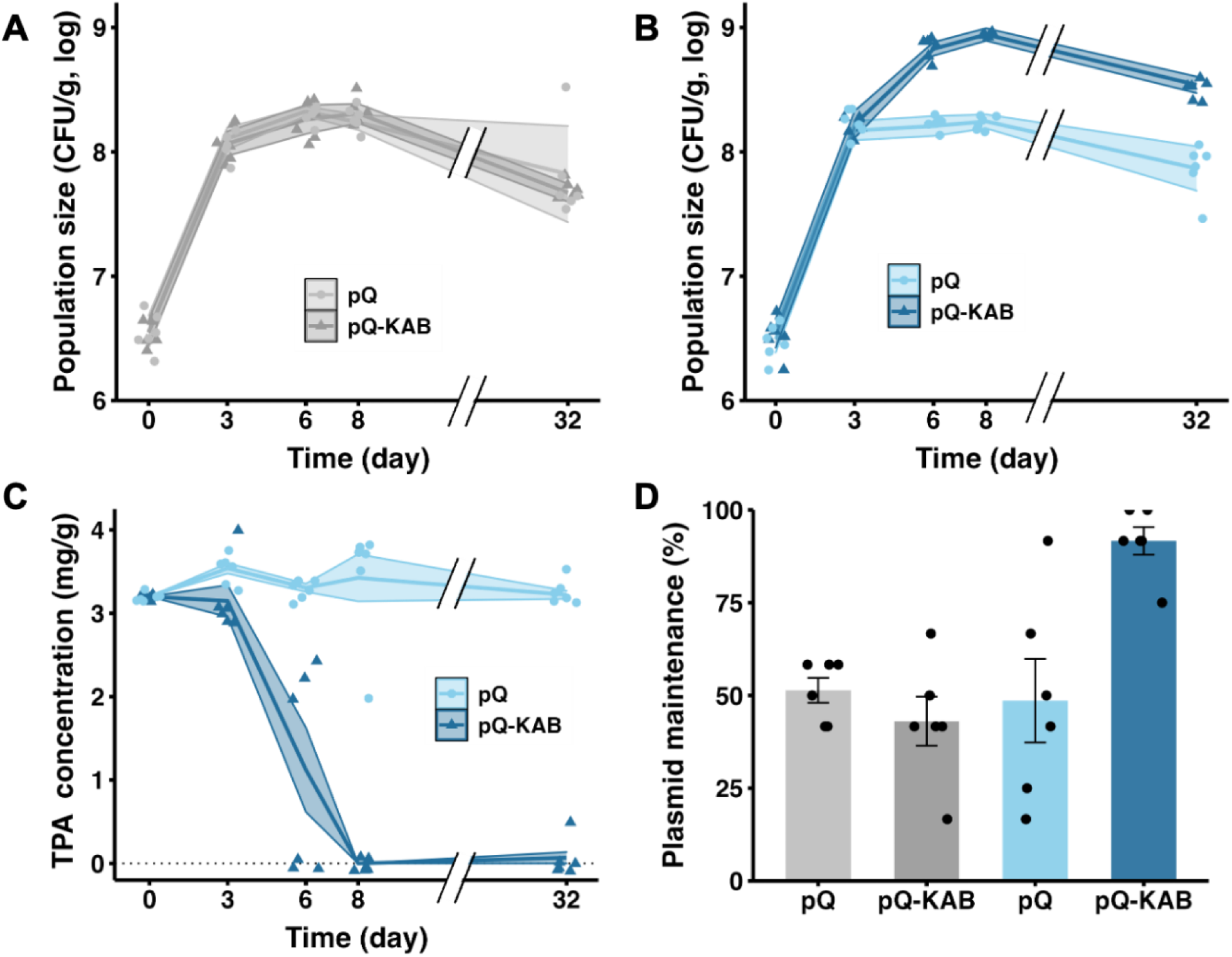
(**A**) Population density of pQBR57 and pQBR57-KAB bearing *P. putida* in soil microcosms without TPA supplementation. (**B**) Identical to panel A with the exception of TPA-supplemented soil microcosms. (**C**) Degradation of TPA in soil microcosms inoculated with pQBR57 and pQBR57-KAB bearing *P. putida.* (**D**) Plasmid maintenance after 32-day incubation for both plasmid genotypes without (grey colors) and with TPA supplementation (blue colors).

### Plasmid transfer in soil microcosm and transconjugant bioremediation function

Interspecies conjugative transfer of pQBR57 or pQBR57-KAB from *P. putida* to *P. fluorescens* was assessed in soil microcosms with or without 3.2 mg/g TPA. The effect of TPA significantly affected transconjugant count while the effect of plasmid genotype was not significant (**Figure 5A**, two-way ANOVA, effect of TPA, F_1,20_ = 84.40, p < 0.001; effect of plasmid, F_1,20_ = 1.25, p = 0.2). *P. fluorescens* transconjugants with either pQBR57 or pQBR57-KAB were assessed for their ability to degrade and utilize TPA in M9 media with 10 mM TPA as the sole carbon source. After 180-hours of incubation, none of the pQBR57 transconjugants could either grow or degrade TPA, whereas 45% (9/20) of *P. fluorescens* pQBR57-KAB transconjugants reached stationary phase by consuming the TPA (**Figure 5C**, Wilcoxon test, W = 36, p = 0.002).

**Figure 5.**
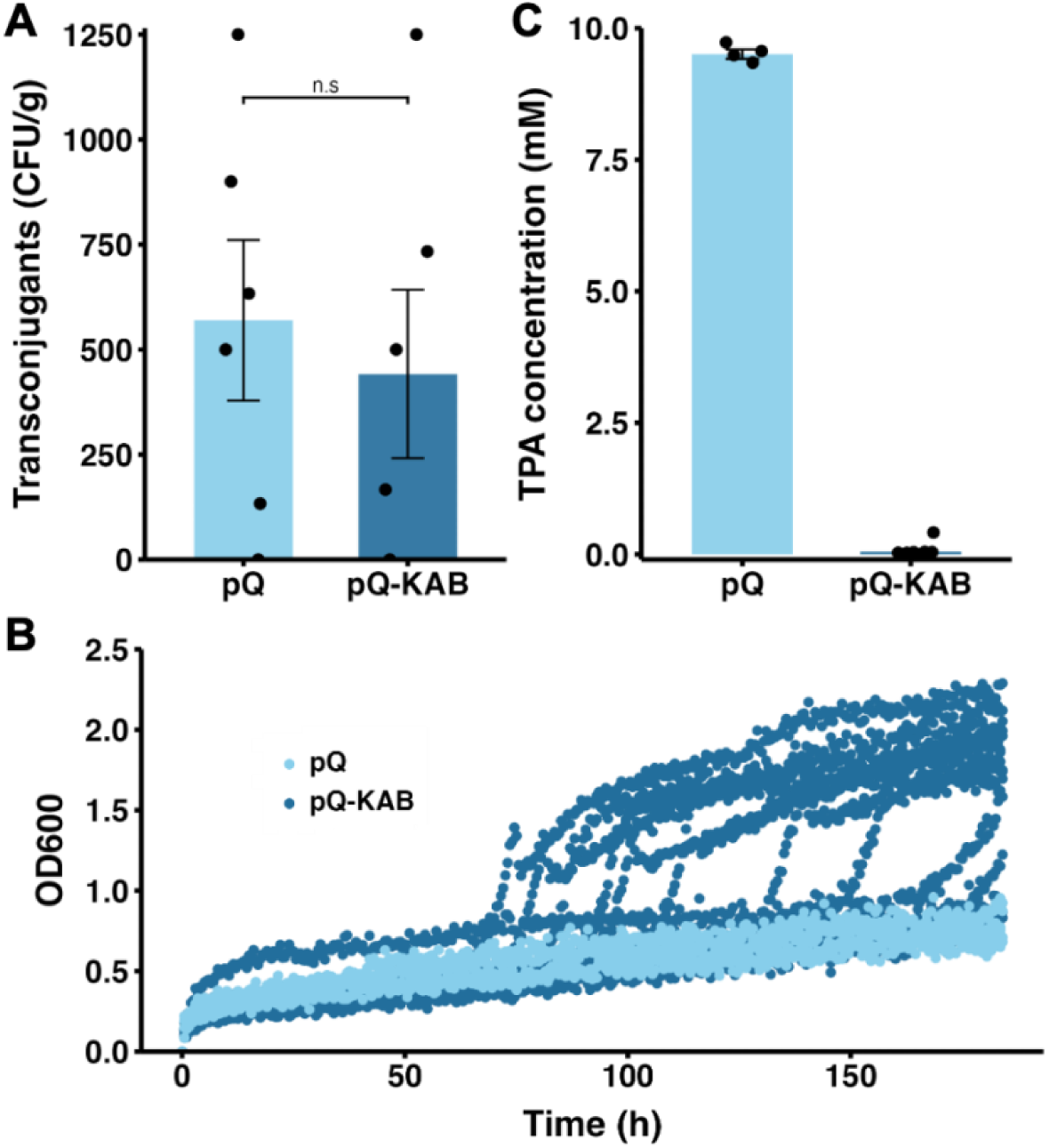
(**A**) Interspecies plasmid transfer in TPA-supplemented soil microcosms with *P. putida* as donor and *P. fluorescens* as recipient. (**B**) Growth dynamics of *P. fluorescens* transconjugants of pQBR57 and pQBR57-KAB in minimal media with 10 mM TPA. (**C**) Final TPA concentration after growth dynamics experiment.

## Discussion

TPA is an environmental pollutant associated with PET plastic manufacture and breakdown. Here, we have built a genetic bioaugmentation vector using a synthetic catabolic operon (KAB) and an environmental plasmid (pQBR57), and further demonstrated the bioremediation function, maintenance, and interspecies transfer of this plasmid in TPA-contaminated soil microcosms. Our findings support the utility of genetic bioaugmentation using engineered environmental plasmids for *in situ* genetic engineering of soil microbiomes to enhance their bioremediation potential. While the use of environmental plasmids for genetic bioaugmentation has been previously reported^28^, most studies are limited to the native catabolic operons found naturally in environmental conjugative plasmids^55^. Expanding the function of environmental plasmids with new bioremediation properties via genetic engineering could greatly expand the range of pollutants to target, for instance, Ke *et al* (2022) knocked-in an amidase leading to expansion the catabolic substrate range of plasmid pDCA-1^56^. However, integration of a complete catabolic operon to equip an environmental plasmid with bioremediation properties is, to the best of our knowledge, reported for the first time in this study.

Genetic bioaugmentation relies on two pillars: (i) transfer and (ii) transconjugant expression of the catabolic payload^57^, which were both experimentally validated in this study. The transfer of pQBR57-KAB from *P. putida* to *P. fluorescens* was demonstrated in soil (**Figure 5A**). *P. fluorescens* is a highly proficient plasmid host capable of stably maintaining and transferring conjugative plasmids to indigenous soil communities^58,59^ acting as a hub for HGT. As such, *P. fluorescens* could drive higher rates of plasmid spread within complex soil communities via secondary transfer, which has been previously reported for other microbiome engineering plasmids^31^. Transconjugant expression of the catabolic payload was validated by the ability of *P. fluorescens* pQBR57-KAB to degrade and utilize TPA in minimal media (**Figure 5B** and **5C**). Consumption of TPA by *P. fluorescens* has not been reported previously, and as such our findings demonstrate that gaining pQBR57-KAB can convert previously non-bioremediating species into bioremediating ones. Unexpectedly, *P. fluorescens* pQBR57-KAB transconjugants varied markedly in their ability to immediately degrade TPA, but the mechanism for this variation is yet unknown. All tested isolates were confirmed to be carrying the plasmid by PCR (**Supplementary Figure 6**), suggesting that plasmid loss cannot explain the observed variation. One possibility is that *P. fluorescens* pQBR57-KAB transconjugants required physiological or evolutionary adaptation to enable them to utilise TPA for growth. This hypothesis is potentially supported by the extended lag phase observed in our transconjugant growth assays, which is substantially longer and more variable for *P. fluorescens* than for *P. putida* (**Supplementary Figure 7**). It is possible that supplementation of the media with an additional carbon source could have accelerated the gain of TPA degradation, as observed in a previous study focused on toluene degradation by *P. fluorescens*^57^.

The plasmid pQBR57 was isolated from soil^46^ and is known to have a relatively low fitness cost^49^. Addition of the KAB operon to pQBR57 had no effect on the fitness cost of carrying the plasmid in the absence of TPA (**Figure 3B**), suggesting that encoding and expressing this additional genetic material did not comprise a substantial bioenergetic burden for host cells. Crucially, however, the KAB operon conferred a large fitness benefit in the presence of TPA, enabling plasmid carriers to use this carbon source for growth and biomass production, leading to higher total bacterial population densities in TPA-contaminated soil (**Figure 4B**) alongside complete bioremediation of TPA from the contaminated soil substrate (**Figure 4C**). Accordingly, the pQBR57-KAB plasmid was maintained at higher frequencies in TPA-contaminated soil (**Figure 4D**). The combination of no additional fitness cost without TPA and high fitness benefits with TPA suggest that pQBR57-KAB is a promising genetic bioaugmentation system for use in complex natural environments, such as soil. By contrast, we observed a lower interspecies conjugation rate with the addition of the KAB operon (**Figure 3A**), which may indicate barriers to acquisition of plasmids containing the KAB operon that manifest only in *P. fluorescens* but not in *P. putida* recipients. Nevertheless, this effect was not apparent in soil, where the difference in transconjugants count did not differ between pQBR57 and pQBR57-KAB (**Figure 5A**). The increase in plasmid transfer in the presence of TPA is similar to increased transfer previously reported for other bioaugmentation plasmids across a range of pollutants whose degradation provides plasmid carriers with a “privatised” nutrient (e.g. 2,4-dichlorophenoxyacetic acid^23,68^, toluene^57^). Notably, increased transfer with selection is not observed for plasmids that carry resistance traits, such as mercury resistance, due to the selective agent killing potential recipients and thus negating opportunities for conjugation^61^. Variants of pQBR57 with higher than wild-type conjugation rates without higher costs have been experimentally evolved by Kottara *et al* (2016)^62^, and could potentially be employed to boost the efficiency of genetic bioaugmentation. Alternatively, further genetically engineering pQBR57-KAB to add genes that facilitate cell-to-cell contact such as adhesins^63^ or are involved in biofilm formation^64,65^ could increase cell-cell contacts and potentially boost conjugation rate.

Genetic bioaugmentation vectors must carefully balance their catabolic efficiency against the potential costs of expressing heterologous traits in naïve hosts, which could limit their spread. Here, we used a medium strength constitutive promoter (*Pem7*) to control the KAB operon on a low copy number plasmid (1-2 copies per cell), which provided appreciable rates of TPA catabolism at a negligible fitness cost in *P. putida*. Specifically, *P. putida* pQBR57-KAB achieved complete depletion of 10 mM TPA after 100-hours of incubation in liquid media (**Figure 2C**) and 3.2 mg/g TPA after 8 days in soil microcosms (**Figure 4C**). More rapid depletion of TPA has been reported with other genetic systems in other studies. Werner *et al* (2021)^51^ placed a chromosomal KAB operon under the control of a strong constitutive promoter (*Ptac*) in *P. putida*, achieving complete degradation of 45 mM TPA within 36 hours in liquid media. In addition, Alvarez *et al* (2024)^66^ used a high-copy number plasmid encoding the KAB operon regulated by a protocatechuate-inducible promoter in *P. putida*, achieving complete depletion of 10mM TPA within 24 hours in liquid media. Importantly, none of these previous studies assessed the fitness costs of high-level KAB expression nor tested TPA catabolism in the context of bioremediation in a more complex and relevant environment, as demonstrated here for soil. It is highly likely that high levels of expression of catabolic genes will boost bioremediation at the expense of incurring high fitness costs, which could limit the spread of the function in communities by reducing both donor and transconjugant fitness causing plasmid carriers to be more rapidly outcompeted. At the other end of the spectrum, plasmids with low expression of the catabolic payload (minimal fitness cost) can be stably maintained and result in long-term bioremediation rates^22^. Genetic bioaugmentation strategies must balance the speed of bioremediation and the fitness cost in diverse natural communities. In nature, catabolic plasmids may preferentially conjugate into and be maintained by slow-growing microorganisms^67^ which might indicate that low-expression constructs could be preferred in the long run.

Our study demonstrates the utility of engineering environmental plasmids with synthetic catabolic operons for genetic bioaugmentation to achieve efficient bioremediation within complex natural substrates. Release of genetically modified strains and plasmids is tightly regulated at present, limiting the use of genetic bioaugmentation approaches reliant upon genetically engineered plasmids, to contained-use applications. Notably, we observed substantially higher rates of maintenance and interspecies conjugation of pQBR57-KAB in TPA-contaminated soil, suggesting that the plasmid’s long-term persistence may be limited once a site is fully bioremediated, although these long-term dynamics require further study. A potentially promising route to application in the short-term may be to deploy pQBR57-KAB for bioremediation in closed systems that ensure biocontainment, such as bioremediation of materials *ex situ* (i.e., where contaminated material is removed for bioremediation off-site) or in self-contained wastewater bioreactors, where the requisite removal of viable GM strains can subsequently be performed. Engineered environmental plasmids could therefore play an important role in reducing pollution by xenobiotics released during manufacturing and waste processing.

## Authors′ contributions

AMA, KM, LC, and MC, planned and performed the experiments. AMA, MB, and ND analysed the data and wrote the manuscript. All authors read and approved the final version of the manuscript.

## Data availability

All data generated or analysed in this manuscript is available as supplementary data files. Any additional data can be made available from the authors upon reasonable request. Materials can be made available upon reasonable request under a material transfer agreement.

## Materials & Methods

### Vector design and cloning

The vector backbone pQBR57 was obtained from Lilley *et al* (1994)^46^ while the engineered catabolic operon tphKAB from Alvarez *et al* (2024)^65^ and the promoter *Pem7* from pSEVA backbone Garcia-Gutierrez *et al* (2020)^53^. The genetic parts were amplified via PCR using Q5 HF Polymerase 2X (New England Biolabs) and the general method: 95°C for 3 minutes, 30 cycles of 20 s at 95°C, 20 s at 57°C and the required time based on 30 kb/s at 72°C, 5 min at 72°C followed the final cycle. All primers were synthesized by IDT. Gibson assembly with HiFi Assembly Mix (New England Biolabs) was used to clone the constitutive promoter, engineered operon and backbone suicide vector into *E. coli* DH5α. Sanger sequencing, through Eurofins, was used to verify successful insertion of promoter and operon. The construct pTS-1-Pem7_tphKAB was isolated using QiaPrep (Qiagen) and transformed by heat-shock into *E. coli* S-17pir, a conjugation competent strain.

The construct was conjugated into *P. fluorescens* SBW25 harbouring pQBR57 via biparental conjugation. Briefly summarised, *E. coli* S-17pir pTS-1_Pem7-tphKAB and *P. fluorescens* SBW25 pQBR57 were grown at 37°C and 28°C, 180rpm for 24 and 48 hours in Luria Broth media supplemented with 12.5 μg/ml tetracycline and Hg (II) 20 μM, respectively. The strains were washed with 1X M9 salts, mixed at a 1:3 donor-to-recipient volumetric ratio and 30μl of the mixture was dropped in centre of an LB agar plate. After incubation at 28°C for 16 hours, the biofilm was scrapped off, resuspended in 1X M9 salts and plated in dual-antibiotic selection plates - 100 μg/ml tetracycline and Hg (II) 20 μM. After incubation for 48 hours at 28°C, transconjugant colonies were selected and streak on LB agar supplemented with 100 μg/ml tetracycline to ensure single-cross overs, i.e. colonies that have integrated the suicide vector into pQBR57. After incubation for 24 hours at 28°C, successful single-cross over colonies were isolated and inoculated in 250ml conical flask with 50ml LB media, the cultures were grown overnight at 28°C, 180 rpm until stationary phase (OD=1.5). The cultures were plated on LB agar supplemented with 20% sucrose and Hg (II) 20 μM. In the event of double crossover, the loss of the suicide vector backbone which carries a levansucrase enzyme (*sacB*) results in unhindered growth at 20% sucrose. Therefore, double-cross over colonies grow larger colonies enabling their selection. Verification of double-cross over was achieved via colony PCR using Phire Hot Start (Thermo Fisher) and the following protocol: 95°C for 3 minutes, 30 cycles of 10 s at 95°C, 10 s at 57°C and the required time based on 10 kb/s at 72°C, 2 min at 72°C followed the final cycle.

Once assembled, the construct pQBR57 Pem7-tphKAB (pQBR57-KAB) and the unmodified pQBR57 were conjugated into *P. putida* KT2440, carrying a streptomycin resistance cassette for selection, from *P. fluorescens* SBW25 via conjugation. Briefly summarised, both strains were grown in 5ml LB overnight at 28°C, 180rpm. The strains were mixed at a 1:3 donor-to-recipient and re-inoculated into fresh 5ml LB at a 1/100 dilution. After overnight incubation at 28°C, 180rpm, plating on dual-antibiotic (100 μg/ml streptomycin and Hg (II) 20 μM) ensured the selection of *P. putida* KT2440 transconjugants.

### Bioremediation function of P. putida pQBR57-KAB

Disodium terephthalate (Thermo Fisher) was dissolved in milli-Q water at a concentration of 500mM via vigorous vortexing for 10 minutes at room temperature. M9 minimal media was supplemented with 10 mM disodium terephthalate as the sole carbon source to make M9 + 10 mM TPA. Three biological replicates, with 4 technical replicates each, of *P. putida* KT2440 pQBR57-KAB were grown in LB, supplemented with Hg (II) 20 μM to ensure plasmid maintenance, for 16 hours at 28°C and 180rpm. After washing with 1X M9 salts, the cultures were diluted 1/100 in M9 + 10 mM TPA and incubated at 28°C in the Biolector for 160 hours. At the end of the experiments, 500 μl of culture was spun at 16G, 5 minutes, 300 μl of supernatant was diluted with milli-Q to the detectable TPA range (0 – 0.5 mM TPA) (**Supplementary Figure 2**) and mixed with 300 μl of methanol and 13 μl of 6M HCl. The mixture was thoroughly mixed and incubated at room temperature for 5 min, followed by centrifugation at 16G for 8 minutes. 500 μl of the resulting supernatant was loaded into High Pressure Liquid Chromatography (HPLC).

To test the bioremediation activity in soil microcosms, 10 grams of John Innes No. 2 compost soil were added to a 30ml glass microcosm and autoclaved twice. Soil microcosms were inoculated with 305μl of 500mM disodium terephthalate solution for a total of 3.2 mg TPA per gram of soil. Overnight cultures, grown in King’s Broth (KB) media, of 6 independent transconjugant lines were washed in1X M9 salts and an aliquot of 100 μl was added to each microcosm, followed by 30-second vortexing at maximum speed. The microcosms were incubated at 28°C and 80% humidity in a static incubator for 3, 6, 8 and 32 days.

### Soil microcosms sample preparation

To determine colony forming units (CFU) and concentration of TPA, 10 ml of milli-Q water and 15 autoclaved glass beads were added to each microcosm, followed by 30-second vortexing at maximum speed. After 5 minutes incubation at room temperature to allow the soil to set, 1 ml of soil wash was extracted. To determine CFU counts, serial dilutions of soil wash were plated in King’s Broth (KB) agar plates and incubated for 24 hours at 28°C. To determine TPA concentration, soil wash was centrifuge at 16G, 5 minutes; 300μl of supernatant was diluted with milli-Q to the detectable TPA range (0 – 0.5 mM) (**Supplementary Figure 2**) and then mixed with 300μl of methanol and 13μl of 6M HCl. The mixture was incubated at room temperature for 5 minutes and centrifuge at 16G, 8min; 500μl of supernatant was added to HPLC tubes and taken to the HPLC.

### High Pressure Liquid Chromatography detection of TPA

Agilent 1200 Infinity Series HPLC instrument equipped with a diode array detector was. The stationary phase was a Kinetex 5 μm C18 100 A column, 250 x 4.6 mm. The mobile phase was a linear gradient of acetonitrile-to-water (5% : 95%) to (30% : 70%) over 20 minutes with 4 minutes post run at (5% : 95%) to re-equilibrate the column before the next injection. All solvents contained 0.1% TFA, the flow rate was 0.8 mL per minute and the sample volume injected per run was 10 μL.

### Relative fitness using flow cytometry

Six colonies of reference (*P. putida* KT2440 pQBR57) and test (*P. putida* KT2440.YPF pQBR57-KAB) strains were grown in 5ml of KB media, supplemented with Hg (II) 20 μM for the plasmid-bearing strains, at 28°C 180rpm for 16 hours. After one wash step with M9 1X Salts, 25 μl of reference and 25 μl of test strains were mixed and inoculated into 5ml of the corresponding media composition. In parallel, a control competition to account for the fitness cost of YFP-expression was set up between *P. putida* KT2440 and *P. putida* KT2440.YFP. To create the matrix of media, 10% Glucose solution and 0.1M disodium terephthalate solutions were diluted accordingly into M9 media. The strains were co-cultured for 96 hours at 28°C 180rpm. Daily samples of 100μl each were prepared for flow cytometry analysis (Cytoflex). Initially, samples were diluted to OD < 0.1 and incubated with 5μg/ml of Hoechst 34580 (ThermoFischer) in the dark for at least 30 minutes. The Cytoflex protocol consisted of 20μl/min flowrate, a gated stop function at 30,000 counts at blue fluorescence above 10^3.5^ and side scattering above 10^3^. In between each sample well, a blank well was run for 40 seconds to detect any possible carry-over.

Rstudio package flowCore was used to curate and analyse, and ggplot2 to plot the data. The final selection rate results from subtracting the selection rate of the control competition (*P. putida* KT2440 vs. *P. putida* KT2440.YFP) to the selection rate of the test competition (*P. putida* KT2440 pQBR57 vs. *P. putida* KT2440.YFP pQBR57-KAB)

### Conjugation rate in liquid media

Donor strains harbour a genomic streptomycin resistance and recipient strains with a genomic gentamycin resistance to aid with selection and identification of transconjugants. Six colonies of donor and recipient were grown on KB media at 28°C 180rpm for 16 hours, supplemented with Hg (II) 20 μM for donor. After one wash step with KB to remove Hg (II) 20 μM from donor cultures, 25μl of donor and 25μl of recipient were mixed, inoculated into 5ml of KB media and grown for 24 hours at 28°C 180rpm. Plating on streptomycin (100μg/ml) yields donor count, on gentamycin (10μg/ml) recipient count and on dual selection with gentamycin and Hg (II) 20 μM yields transconjugant count. After incubation at 28°C for 24 hours and colony counting, the conjugation rate was calculated using Simonsen method^54^.

### Plasmid transfer in soil microcosm and transconjugant bioremediation function

To test the plasmid transfer in soil, soil microcosms were prepared as previously stated. Six colonies of donor (*P. putida*) for each plasmid, pQBR57 and pQBR57-KAB, and recipients, either *P. putida* (intraspecies) or *P. fluorescens* (interspecies), were grown on KB media at 28°C 180rpm for 16 hours, supplemented with Hg (II) 20 μM for the plasmid carriers. After one wash step with KB to remove the mercury from donor cultures, 25μl aliquots of each the donor and recipient were mixed and inoculated into soil microcosms. The microcosms were incubated in a static incubator at 28°C and 80% humidity. These were sampled as previously stated after 24 and 72 hours. To determine the presence of transconjugants, dual selection of gentamycin (10 μg/ml) and Hg (II) 20μM was used. Incubation at 28°C for 24 hours resulted in transconjugant colonies. To confirm the bioremediation phenotype of these soil transconjugants, they were grown on KB supplemented with Hg (II) 20μM at 28°C, 180rpm for 16 hours, then washed with1X M9 Salts to remove any carry-over nutrients and diluted 1/100 into M9 + 10 mM TPA in a 48-well flower plate. The growth on TPA was recorded for 160 hours in the Biolector and endpoint sampling was used to determine the concentration of TPA after the incubation period using HPLC, as previously explained.

## Supplementary Information

**Supplementary Figure 1.**
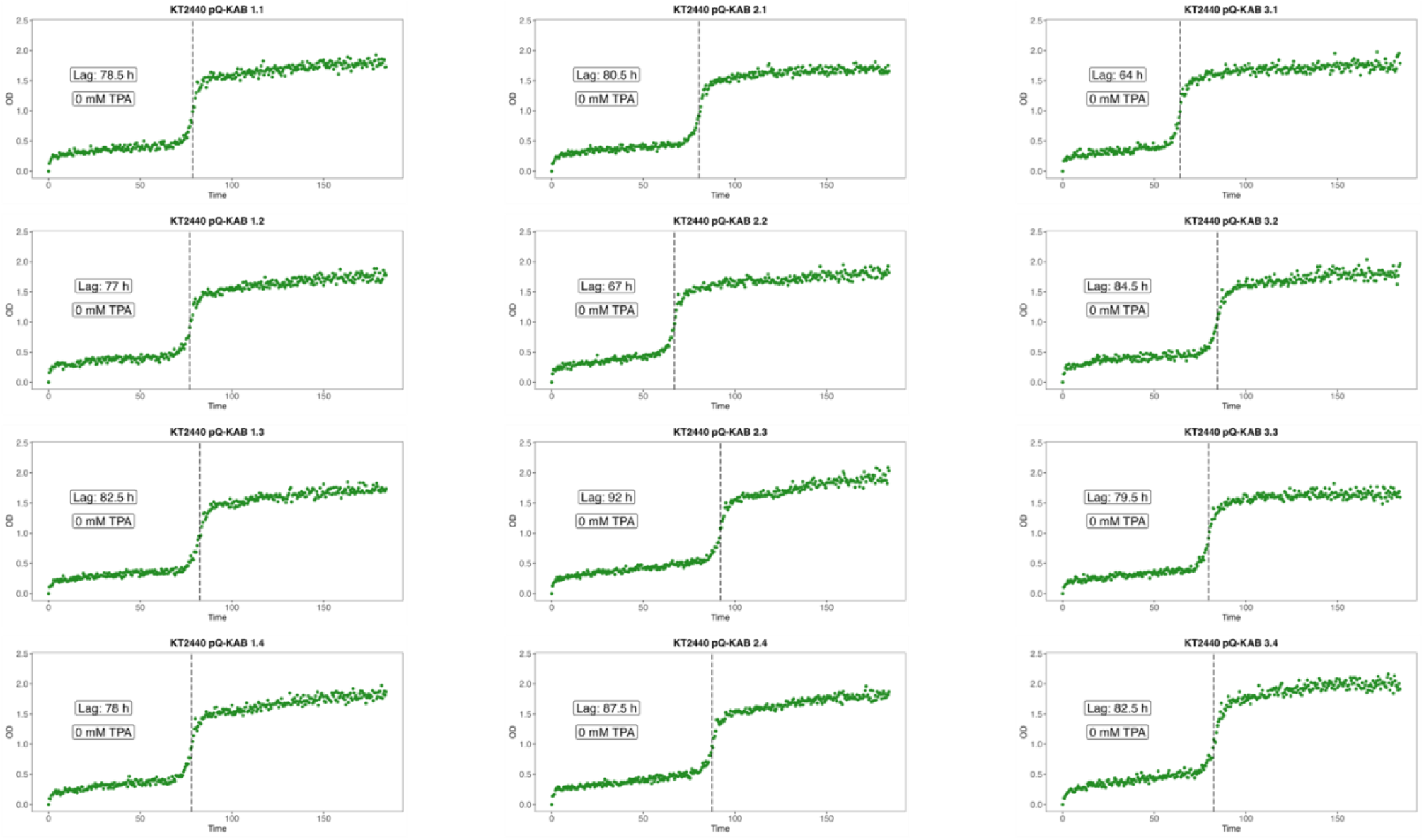
*P. putida* pQBR57-KAB each individual replicate growth curves, with TPA as the sole carbon source, the lag-phase duration and final TPA concentration indicated.

**Supplementary Figure 2.**
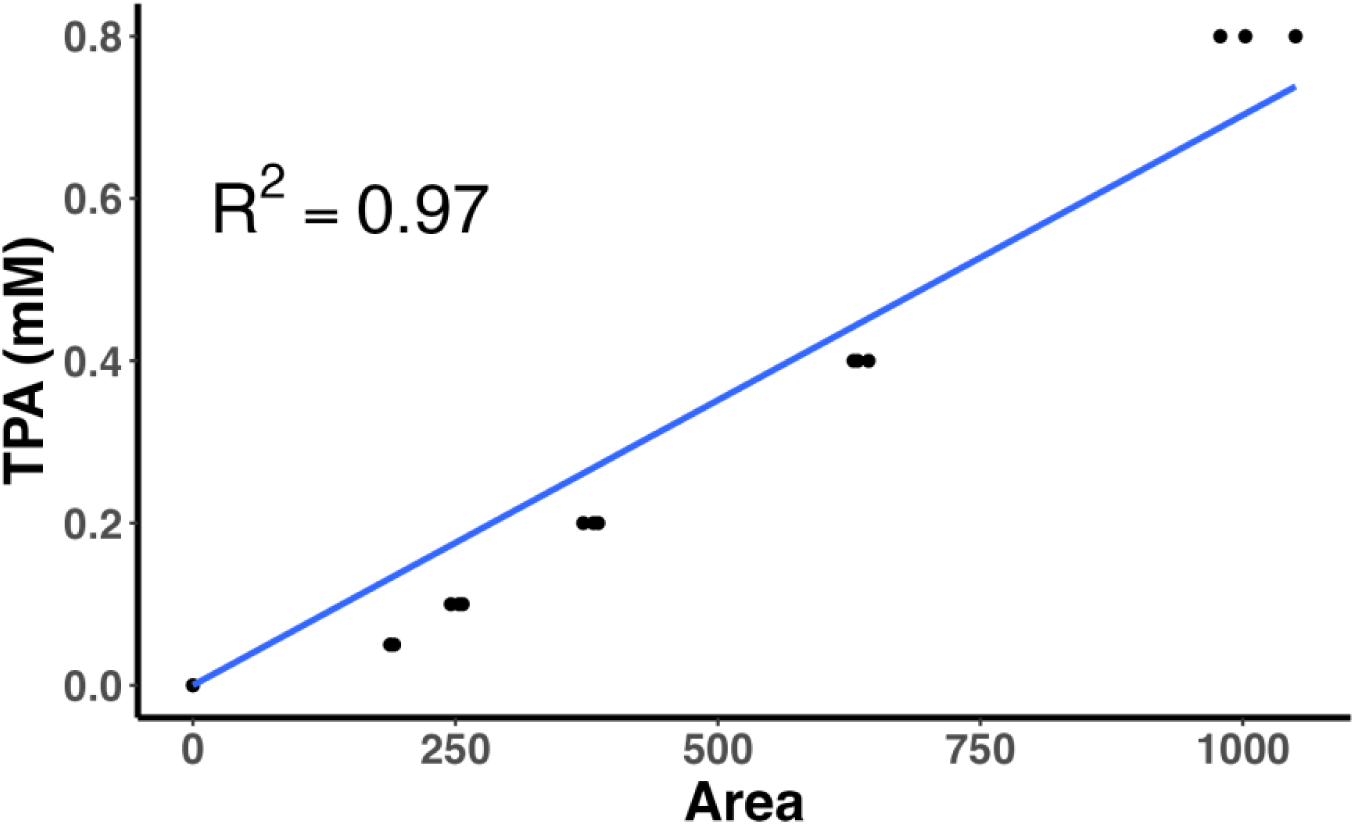
TPA standard curve for HPLC.

**Supplementary Figure 3.**
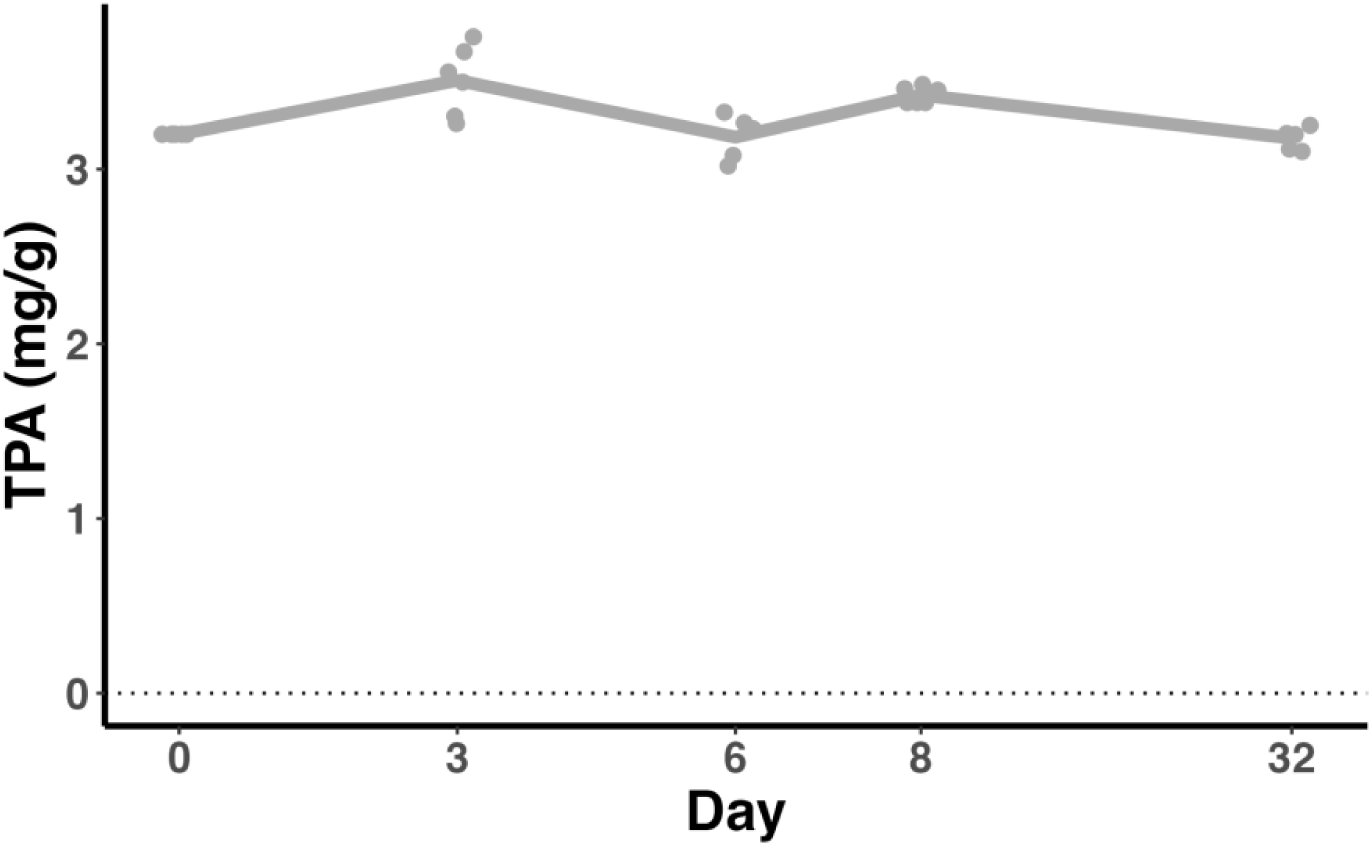
Control of TPA degradation in soil, no bacteria were inoculated.

**Supplementary Figure 4.**
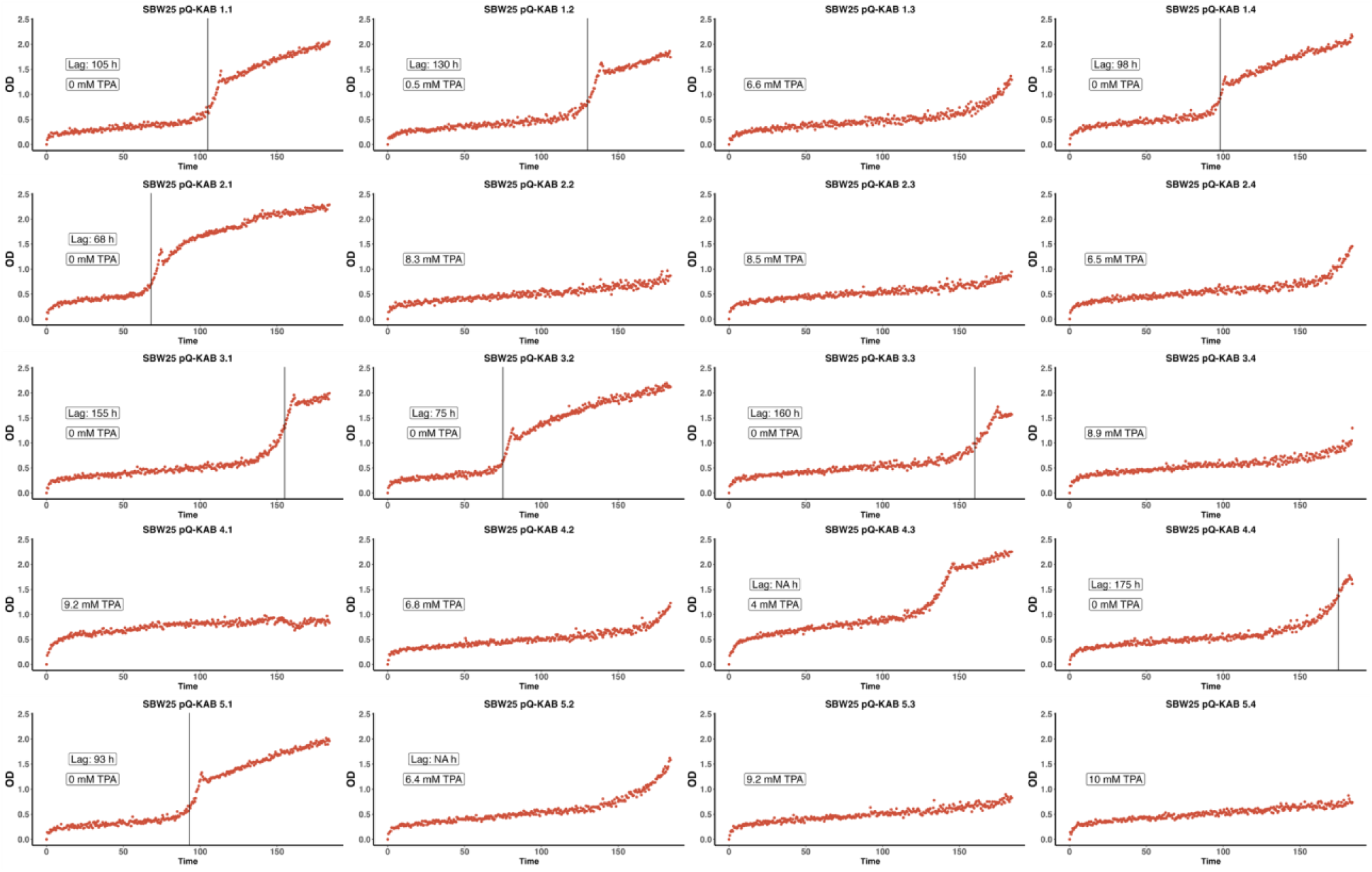
Soil-isolated transconjugants of *P. fluorescens* pQBR57-KAB growing on M9 + 10mM TPA. The final concentration of TPA and the duration of the lag-phase are included in each plot.

**Supplementary Figure 5.**
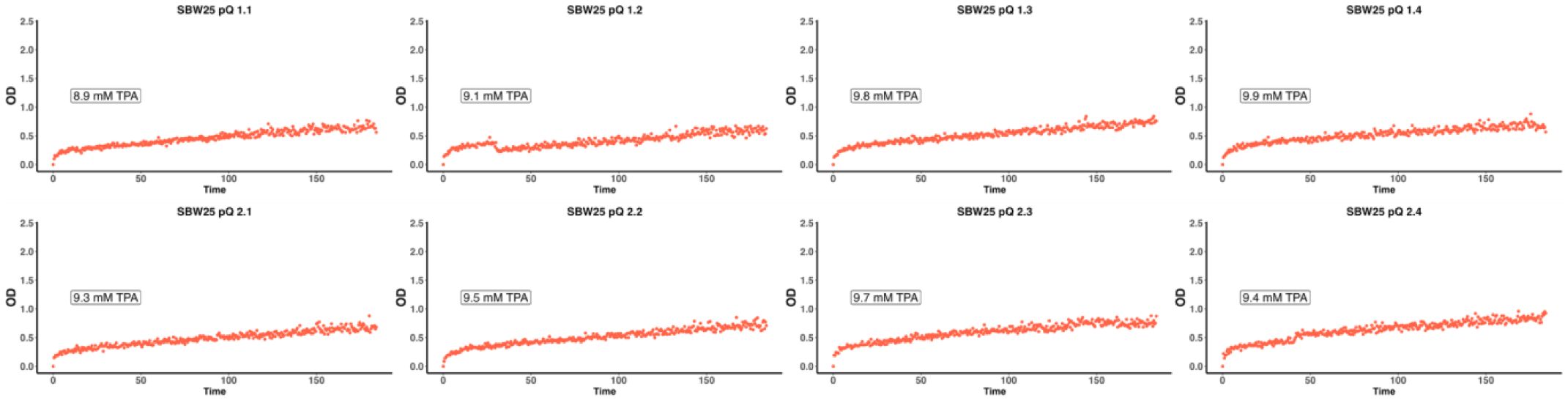
Soil-isolated transconjugants of *P. fluorescens* pQBR57 growing on M9 + 10mM TPA. The final concentration of TPA is included in each plot.

**Supplementary Figure 6.**
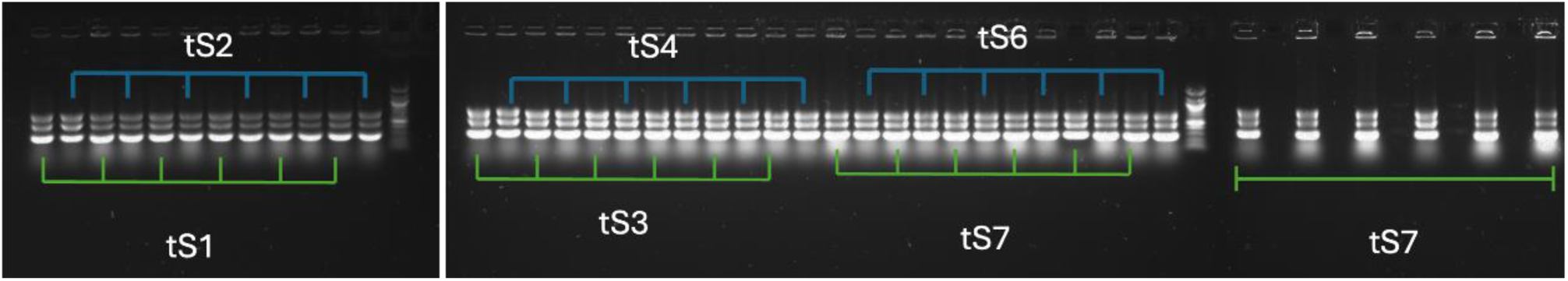
PCR confirmation of soil-isolated transconjugants of *P. fluorescens* pQBR57-KAB. The reaction is multiplexed and its targeting *merA*, *tphK* and *uvrD* in pQBR57-KAB.

**Supplementary Figure 7.**
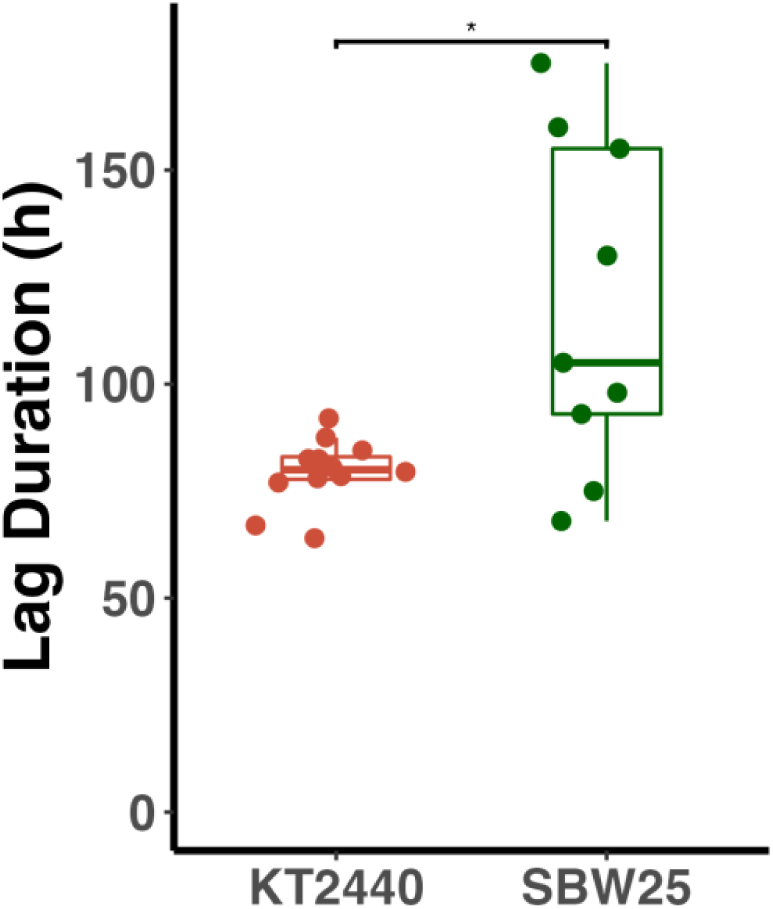
Difference in lag duration of *P. putida* and *P. fluorescens* carrying pQBR57-KAB growing on M9 + 10 mM TPA.

## References

1. Urbanek, A. K., Kosiorowska, K. E. & Mirończuk, A. M. Current Knowledge on Polyethylene Terephthalate Degradation by Genetically Modified Microorganisms. Front. Bioeng. Biotechnol. 9, 771133 (2021).

2. De Vos, L., Van de Voorde, B., Van Daele, L., Dubruel, P. & Van Vlierberghe, S. Poly(alkylene terephthalate)s: From current developments in synthetic strategies towards applications. Eur. Polym. J. 161, 110840 (2021).

3. Pophali, G. R., Khan, R., Dhodapkar, R. S., Nandy, T. & Devotta, S. Anaerobic–aerobic treatment of purified terephthalic acid (PTA) effluent; a techno-economic alternative to two-stage aerobic process. J. Environ. Manage. 85, 1024–1033 (2007).

4. Verma, S., Prasad, B. & Mishra, I. M. Treatment of Petrochemical Wastewater by Acid Precipitation and Carbon Adsorption. J. Hazard. Toxic Radioact. Waste 18, 04014013 (2014).

5. Lee, Y.-S. & Han, G.-B. Treatment of Wastewater from Purified Terephtalic Acid (PTA) Production in a Two-stage Anaerobic Expanded Granular Sludge Bed System. Environ. Eng. Res. 19, 355–361 (2014).

6. Ball, G. L., McLellan, C. J. & Bhat, V. S. Toxicological review and oral risk assessment of terephthalic acid (TPA) and its esters: A category approach. Crit. Rev. Toxicol. 42, 28–67 (2012).

7. Daramola, M. O., Aransiola, E. F. & Adeogun, A. G. Comparative study of thermophilic and mesophilic anaerobic treatment of purified terephthalic acid (PTA) wastewater. Nat. Sci. 03, 371 (2011).

8. Goel, R., Jayal, P., Negi, H., Saravanan, P. r. & Zaidi, M. g. h. Comparative in situ PET biodegradation assay using indigenously developed consortia. Int. J. Environ. Waste Manag. 13, 348–361 (2014).

9. Farzi, A., Dehnad, A. & Fotouhi, A. F. Biodegradation of polyethylene terephthalate waste using *Streptomyces* species and kinetic modeling of the process. Biocatal. Agric. Biotechnol. 17, 25–31 (2019).

10. Roberts, C. et al. Environmental Consortium Containing Pseudomonas and Bacillus Species Synergistically Degrades Polyethylene Terephthalate Plastic. mSphere 5, (2020).

11. Yoshida, S. et al. A bacterium that degrades and assimilates poly(ethylene terephthalate). Science 351, 1196–1199 (2016).

12. Sasoh, M. et al. Characterization of the Terephthalate Degradation Genes of Comamonas sp. Strain E6. Appl. Environ. Microbiol. 72, 1825–1832 (2006).

13. Hara, H., Eltis, L. D., Davies, J. E. & Mohn, W. W. Transcriptomic analysis reveals a bifurcated terephthalate degradation pathway in Rhodococcus sp. strain RHA1. J. Bacteriol. 189, 1641–1647 (2007).

14. Narancic, T. et al. Genome analysis of the metabolically versatile Pseudomonas umsongensis GO16: the genetic basis for PET monomer upcycling into polyhydroxyalkanoates. Microb. Biotechnol. 14, 2463–2480 (2021).

15. Gautom, T. et al. Structural basis of terephthalate recognition by solute binding protein TphC. Nat. Commun. 12, 6244 (2021).

16. Salvador, M. et al. Microbial Genes for a Circular and Sustainable Bio-PET Economy. Genes 10, 373 (2019).

17. Li, H. et al. Improved anaerobic degradation of purified terephthalic acid wastewater by adding nanoparticles or co-substrates to facilitate the electron transfer process. Environ. Sci. Nano 9, 1011–1024 (2022).

18. Wen, A. et al. Enabling Biological Nitrogen Fixation for Cereal Crops in Fertilized Fields. ACS Synth. Biol. 10, 3264– 3277 (2021).

19. Brophy, J. A. N. et al. mini ICEB1. Nat. Microbiol. 3, 1043–1053 (2018).

20. Pang, F. et al. Soil phosphorus transformation and plant uptake driven by phosphate-solubilizing microorganisms. Front. Microbiol. 15, (2024).

21. Vassileva, M. et al. Multifunctional properties of phosphate-solubilizing microorganisms grown on agro-industrial wastes in fermentation and soil conditions. Appl. Microbiol. Biotechnol. 85, 1287–1299 (2010).

22. Gao, C., Jin, X., Ren, J., Fang, H. & Yu, Y. Bioaugmentation of DDT-contaminated soil by dissemination of the catabolic plasmid pDOD. J. Environ. Sci. China 27, 42–50 (2015).

23. Top, E. M., Van Daele, P., De Saeyer, N. & Forney, L. J. Enhancement of 2,4-dichlorophenoxyacetic acid (2,4-D) degradation in soil by dissemination of catabolic plasmids. Antonie Van Leeuwenhoek 73, 87–94 (1998).

24. Ren, C. et al. Genetic Bioaugmentation of Activated Sludge with Dioxin-Catabolic Plasmids Harbored by Rhodococcus sp. Strain p52. Environ. Sci. Technol. 52, 5339–5348 (2018).

25. Mrozik, A. & Piotrowska-Seget, Z. Bioaugmentation as a strategy for cleaning up of soils contaminated with aromatic compounds. Microbiol. Res. 165, 363–375 (2010).

26. Atuchin, V. V. et al. Microorganisms for Bioremediation of Soils Contaminated with Heavy Metals. Microorganisms 11, 864 (2023).

27. Pande, V., Pandey, S. C., Sati, D., Bhatt, P. & Samant, M. Microbial Interventions in Bioremediation of Heavy Metal Contaminants in Agroecosystem. Front. Microbiol. 13, 824084 (2022).

28. Garbisu, C., Garaiyurrebaso, O., Epelde, L., Grohmann, E. & Alkorta, I. Plasmid-Mediated Bioaugmentation for the Bioremediation of Contaminated Soils. Front. Microbiol. 8, 1966 (2017).

29. Khan, I. et al. Mechanism of the Gut Microbiota Colonization Resistance and Enteric Pathogen Infection. Front. Cell. Infect. Microbiol. 11, 716299 (2021).

30. Ramos, J. L., Duque, E. & Ramos-Gonzalez, M. I. Survival in soils of an herbicide-resistant Pseudomonas putida strain bearing a recombinant TOL plasmid. Appl. Environ. Microbiol. 57, 260–266 (1991).

31. Ronda, C., Chen, S. P., Cabral, V., Yaung, S. J. & Wang, H. H. Metagenomic engineering of the mammalian gut microbiome in situ. Nat. Methods 16, 167–170 (2019).

32. De Rore, H. et al. Transfer of the catabolic plasmid RP4::Tn4371 to indigenous soil bacteria and its effect on respiration and biphenyl breakdown. FEMS Microbiol. Ecol. 15, 71–77 (1994).

33. Inoue, D. et al. Impacts of gene bioaugmentation with pJP4-harboring bacteria of 2,4-D-contaminated soil slurry on the indigenous microbial community. Biodegradation 23, 263–276 (2012).

34. de Lipthay, J. R., Barkay, T. & Sørensen, S. J. Enhanced degradation of phenoxyacetic acid in soil by horizontal transfer of the tfdA gene encoding a 2,4-dichlorophenoxyacetic acid dioxygenase. FEMS Microbiol. Ecol. 35, 75–84 (2001).

35. Dejonghe, W. et al. Effect of Dissemination of 2,4-Dichlorophenoxyacetic Acid (2,4-D) Degradation Plasmids on 2,4-D Degradation and on Bacterial Community Structure in Two Different Soil Horizons. Appl. Environ. Microbiol. 66, 3297–3304 (2000).

36. French, K. E., Zhou, Z. & Terry, N. Horizontal ‘gene drives’ harness indigenous bacteria for bioremediation. Sci. Rep. 10, 15091 (2020).

37. Bottery, M. J. Ecological dynamics of plasmid transfer and persistence in microbial communities. Curr. Opin. Microbiol. 68, 102152 (2022).

38. Springael, D. & Top, E. M. Horizontal gene transfer and microbial adaptation to xenobiotics: new types of mobile genetic elements and lessons from ecological studies. Trends Microbiol. 12, 53–58 (2004).

39. Shintani, M. et al. Characterization of the replication, maintenance, and transfer features of the IncP-7 plasmid pCAR1, which carries genes involved in carbazole and dioxin degradation. Appl. Environ. Microbiol. 72, 3206–3216 (2006).

40. Siddavattam, D., Yakkala, H. & Samantarrai, D. Lateral transfer of organophosphate degradation (opd) genes among soil bacteria: mode of transfer and contributions to organismal fitness. J. Genet. 98, 23 (2019).

41. Molbak, L., Licht, T., Kvist, T., Kroer, N. & Andersen, S. Plasmid Transfer from Pseudomonas putida to the Indigenous Bacteria on Alfalfa Sprouts: Characterization, Direct Quantification, and In Situ Location of Transconjugant Cells. Appl. Environ. Microbiol. 69, 5536–42 (2003).

42. Kasai, Y., Inoue, J. & Harayama, S. The TOL Plasmid pWW0 xylN Gene Product from Pseudomonas putida Is Involved in m-Xylene Uptake. J. Bacteriol. 183, 6662–6666 (2001).

43. Ono, A. et al. Isolation and characterization of naphthalene-catabolic genes and plasmids from oil-contaminated soil by using two cultivation-independent approaches. Appl. Microbiol. Biotechnol. 74, 501–510 (2007).

44. Filonov, A. E. et al. Horizontal transfer of catabolic plasmids and naphthalene biodegradation in open soil. Microbiology 79, 184–190 (2010).

45. Jussila, M. M., Zhao, J., Suominen, L. & Lindström, K. TOL plasmid transfer during bacterial conjugation in vitro and rhizoremediation of oil compounds in vivo. Environ. Pollut. Barking Essex 146, 510–524 (2007).

46. Lilley, A. K., Fry, J. C., Day, M. J. & Bailey, M. J. In situ transfer of an exogenously isolated plasmid between Pseudomonas spp. in sugar beet rhizosphere. Microbiology 140, 27–33 (1994).

47. Hall, J. P. J., Harrison, E., Pärnänen, K., Virta, M. & Brockhurst, M. A. The Impact of Mercury Selection and Conjugative Genetic Elements on Community Structure and Resistance Gene Transfer. Front. Microbiol. 11, (2020).

48. Kottara, A., Carrilero, L., Harrison, E., Hall, J. P. J. & Brockhurst, M. A. The dilution effect limits plasmid horizontal transmission in multispecies bacterial communities. Microbiology 167, 001086 (2021).

49. Hall, J. P. J. et al. Environmentally co-occurring mercury resistance plasmids are genetically and phenotypically diverse and confer variable context-dependent fitness effects. Environ. Microbiol. 17, 5008–5022 (2015).

50. Hall, J. P. J. et al. Plasmid fitness costs are caused by specific genetic conflicts enabling resolution by compensatory mutation. PLoS Biol. 19, e3001225 (2021).

51. Werner, A. Z. et al. Tandem chemical deconstruction and biological upcycling of poly(ethylene terephthalate) to β-ketoadipic acid by *Pseudomonas putida* KT2440. Metab. Eng. 67, 250–261 (2021).

52. Kincannon, W. M., et al. Biochemical and structural characterization of an aromatic ring–hydroxylating dioxygenase for terephthalic acid catabolism. Proc. Natl. Acad. Sci. 119, e2121426119 (2022).

53. García-Gutiérrez, C. et al. Multifunctional SEVA shuttle vectors for actinomycetes and Gram-negative bacteria. MicrobiologyOpen 9, 1135–1149 (2020).

54. Simonsen, L., Gordon, D. M., Stewart, F. M. & Levin, B. R. Estimating the rate of plasmid transfer: an end-point method. Microbiology 136, 2319–2325 (1990).

55. Cycoń, M., Mrozik, A. & Piotrowska-Seget, Z. Bioaugmentation as a strategy for the remediation of pesticide-polluted soil: A review. Chemosphere 172, 52–71 (2017).

56. Ke, Z. et al. Engineering of the chloroaniline-catabolic plasmid pDCA-1 and its potential for genetic bioaugmentation. Int. Biodeterior. Biodegrad. 172, 105435 (2022).

57. Ikuma, K. & Gunsch, C. K. Functionality of the TOL plasmid under varying environmental conditions following conjugal transfer. Appl. Microbiol. Biotechnol. 97, 395–408 (2013).

58. Crowley, D. E., Brennerova, M. V., Irwin, C., Brenner, V. & Focht, D. D. Rhizosphere effects on biodegradation of 2,5-dichlorobenzoate by a bioluminescent strain of root-colonizing *Pseudomonas fluorescens*. FEMS Microbiol. Ecol. 20, 79–89 (1996).

59. Sarand, I., Haario, H., Jørgensen, K. S. & Romantschuk, M. Effect of inoculation of a TOL plasmid containing mycorrhizosphere bacterium on development of Scots pine seedlings, their mycorrhizosphere and the microbial flora in m-toluate-amended soil. FEMS Microbiol. Ecol. 31, 127–141 (2000).

60. DiGiovanni, G. D., Neilson, J. W., Pepper, I. L. & Sinclair, N. A. Gene transfer of Alcaligenes eutrophus JMP134 plasmid pJP4 to indigenous soil recipients. Appl. Environ. Microbiol. 62, 2521–2526 (1996).

61. Stevenson, C., Hall, J. P., Harrison, E., Wood, Aj. & Brockhurst, M. A. Gene mobility promotes the spread of resistance in bacterial populations. ISME J. 11, 1930–1932 (2017).

62. Kottara, A., Hall, J. P. J., Harrison, E. & Brockhurst, M. A. Multi-host environments select for host-generalist conjugative plasmids. BMC Evol. Biol. 16, 70 (2016).

63. Robledo, M. et al. Targeted bacterial conjugation mediated by synthetic cell-to-cell adhesions. Nucleic Acids Res. 50, 12938–12950 (2022).

64. Hausner, M. & Wuertz, S. High rates of conjugation in bacterial biofilms as determined by quantitative in situ analysis. Appl. Environ. Microbiol. 65, 3710–3713 (1999).

65. Ghigo, J. M. Natural conjugative plasmids induce bacterial biofilm development. Nature 412, 442–445 (2001).

66. Gonzalez, G. A., Chacόn, M., Berepiki, A., Fisher, K. & Dixon, N. Degradation and bioconversion of complex municipal solid waste streams into human biotherapeutics and biopolymers. Preprint at 10.21203/rs.3.rs-2582698/v1 (2023).

67. Varner, P. M., Allemann, M. N., Michener, J. K. & Gunsch, C. K. The effect of bacterial growth strategies on plasmid transfer and naphthalene degradation for bioremediation. Environ. Technol. Innov. 28, 102910 (2022).

